# Establishment and Characterization of a New Immortalized Human Adenomyosis Epithelial-Like Cell Line, tAEC21

**DOI:** 10.1101/2025.06.16.660025

**Authors:** Yuliya Klymenko, Jessica L. Kersey, Hunter D. Quigley, Shannon M. Hawkins

**Affiliations:** Department of Obstetrics and Gynecology, Indiana University School of Medicine, Indianapolis, IN, United States of America; Department of Biochemistry and Molecular Biology, Indiana University School of Medicine, Indianapolis, IN, United States of America; Indiana University Melvin and Bren Simon Comprehensive Cancer Center, Indiana University School of Medicine, Indianapolis, IN, United States of America

**Keywords:** endometrium, adenomyosis, endometriosis, translational research, immortalization, epithelial cell line, research resource, inflammatory response, three-dimensional, in vitro

## Abstract

Adenomyosis occurs when endometrial glands and stroma grow within the uterine myometrium. Adenomyosis, as a clinically impactful disease, causes significant pelvic pain and heavy menstrual bleeding. Adenomyosis remains understudied due to the paucity of translational research tools and model systems. This study aimed to create a telomerase-transformed epithelial-like cell line derived from the eutopic endometrium of a subject with focal adenomyosis. De-identified endometrium was processed via mechanical and enzymatic digestion. Epithelial and stromal populations were separated via selective adhesion, followed by fluorescence-activated cell sorting with an epithelial cellular adhesion molecule (EpCAM). EpCAM+ cells were efficiently immortalized with the human telomerase reverse transcriptase gene. Analyses confirmed that the cells were human, without mycoplasma contamination, and exhibited a unique 16-marker short tandem repeat profile. Cytogenetic analysis on G-banded metaphase spreads indicated polyploidy with multiple chromosomal rearrangements. The line, designated as tAEC21, expressed epithelial markers cytokeratin-5 and N-cadherin but not stromal marker CD10. Cells avidly responded to tumor necrosis factor-alpha stimulation by upregulating interleukin-6, C-X-C motif chemokine ligand 8, C-C motif chemokine ligand 2, and mucin 1 gene expression. In a heterotypic, three-dimensional spheroid model, tAEC21 displayed a biologically relevant pattern by assembling into an epithelial shell around the stromal cell core. In two-dimensional monolayers, tAEC21 cells were negative for estrogen and progesterone receptors, while as 3D spheroids, tAEC21 exhibited strong positivity for estrogen but not progesterone receptors. This new epithelial-like, adenomyosis-derived cell line, tAEC21, will be an impactful research resource.

## INTRODUCTION

Adenomyosis is a common, benign uterine condition in which endometrial glands and stroma are present within the uterine myometrium. Clinical symptoms of adenomyosis include abnormal uterine bleeding, pelvic pain, and infertility. These symptoms are similar to those of other common gynecologic diseases, including endometrial cancer, endometriosis, primary dysmenorrhea, and uterine leiomyomata. For example, in the evaluation of abnormal uterine bleeding using the PALM-COEIN strategy, adenomyosis is a common structural cause of heavy menstrual and intermenstrual bleeding along with **P**olyps, **A**denomyosis, **L**eiomyomata, uterine **M**alignancy and endometrial hyperplasia [1]. Patients with adenomyosis may have varying degrees of pelvic pain, ranging from intermittent mild pelvic discomfort or abdominal bloating to chronic pelvic pain (defined as more than six months of pelvic pain that affects daily activities), dysmenorrhea (defined as painful menstruation), dyspareunia (defined as painful sexual intercourse), dyschezia (defined as painful defecation), and dysuria (defined as painful urination). Up to 30% of patients present with no symptoms and are diagnosed histologically during hysterectomy for other gynecologic indications [2–5]. Frequent presence of adenomyosis and other gynecological comorbidities with similar symptomatology, in particular, endometriosis (up to 90% concurrency) or uterine leiomyomas (up to 57% concurrency), hinder timely and accurate diagnosis and appropriate intervention – a challenge which is now more effectively overcome with the advanced imaging modalities [6–8]. The definitive pathological criteria for adenomyosis include the presence of endometrial glands and stroma within the myometrium and extending at least 2.5 mm deep from the endometrium-myometrium junction or deeper than 25% of the total myometrial thickness [9]. Histologically, it is divided into diffuse (*i.e.,* multiple ectopic endometrial foci with undefined borders widely scattered throughout the myometrium) or focal (*i.e.,* isolated endometrial gland nodules, localized and well-circumscribed by myometrium) [10].

Furthermore, adenomyosis affects pregnancy rates and obstetrical outcomes. In patients thought to have adenomyosis by imaging findings at the time of infertility evaluation, adenomyosis is associated with lower embryo implantation rates, reduced pregnancy rates, and higher miscarriage rates. The hypothesized pathophysiology involves a hostile uterine environment, impaired adhesion molecule expression, and local inflammation [11, 12]. Data from mouse models of adenomyosis support this hypothesis [13–15]. Previous work has shown that the endometrium from uteri with adenomyosis exhibits a distinct molecular profile that may be responsible for uterine dysfunction [16, 17].

A significant hurdle in adenomyosis research is the scarcity of representative disease models. While patient samples from surgeries are valuable for histological analyses, biomarker discovery, and validating diagnostic tools, they are unsuitable for studying cell behavior, disease progression mechanisms, or testing therapeutic interventions. In that regard, cultured primary epithelial and stromal cells from endometrial or adenomyosis biopsies are preferable. However, primary cultures are challenging to isolate, purify, and culture and have limited scalability compared to immortal cell lines [18, 19]. Meanwhile, in vitro propagation and immortalization of benign endometrial epithelial cells have been tricky [19–21]. In the absence of available immortalized adenomyosis cell lines, researchers mimic endometrium with classic epithelial features by utilizing an epithelial-type Ishikawa human endometrial adenocarcinoma cell line positive for epithelial marker E-cadherin and low/negative for mesenchymal markers of invasiveness N-cadherin and vimentin [22–24]. To model ectopic lesions, a benign endometriotic epithelial-like 12Z cell line with invasive properties (N-cadherin and vimentin-positive and E-cadherin-negative), derived from a red peritoneal endometriotic lesion, is commonly employed [25, 26]. No adenomyosis-derived epithelial cell lines existed until recently. The only published immortalized human adenomyosis ectopic cell line (ihAMEC) was created by simian vacuolating virus 40 (SV40) lentiviral infection and is not commercially available or widely shared [27].

Our work presents a novel adenomyosis-derived epithelial-like cell line immortalized via retroviral transduction of a human telomerase reverse transcriptase (*TERT*) gene. This new cell line retains epithelial morphology, expresses both epithelial and mesenchymal markers (indicating epithelial origin and acquired invasive properties), and exhibits a hormonal receptor profile characteristic of adenomyosis. The cell line is sensitive to immune pressure and retains biologically relevant spatial architecture in a three-dimensional (3D) co-culture model. This cell line offers an additional in vitro research resource to explore the mechanisms of adenomyosis.

## MATERIALS AND METHODS

### Cell Lines

The 12Z cell line was obtained from Applied Biological Materials, Inc. (ABM; #T0764, Vancouver, Canada) and maintained in Dulbecco’s Modified Eagle Medium: Nutrient Mixture F12 (DMEM/F12; #11330-032, Thermo Fisher Scientific, Waltham, MA, USA), supplemented with 10% fetal bovine serum (FBS; #S12450, Atlanta Biologicals, Flowery Branch, GA, USA) and 1% penicillin/streptomycin (P/S; #15140-122, Thermo Fisher Scientific) [25]. The THESC cell line was purchased from American Type Culture Collection (ATCC; #CRL-4003, Manassas, VA, USA) and maintained in phenol red(-) DMEM/F12 medium (#21041-025, Thermo Fisher Scientific), supplemented with 10% charcoal-dextran-treated FBS (#S11650, R&D Systems, Minneapolis, MN, USA), 1% ITS™+Premix Universal Culture Supplement (containing human recombinant insulin, human transferrin, selenous acid, bovine serum albumin, and linoleic acid) (#354352, Corning, Durham, NC, USA), and 500 ng/mL puromycin (#J67236-XF, Thermo Fisher Scientific) [28]. The tAEC21 cells were cultured in human epithelial-stromal [HES, phenol red(-) DMEM/F12 medium + 10% charcoal-dextran treated FBS +1% Anti-Anti + 0.075% sodium bicarbonate (#25080094, Gibco)]. The TOV21G cell line was purchased from ATCC (#CRL-3577, ATCC) and was maintained in a 1:1 ratio of Medium-199 (#M4530, Sigma-Aldrich, St. Louis, MO, USA) and MCBD-105 (#M6395, Sigma-Aldrich) supplemented with 15% FBS and 1% P/S [29]. The MCF-7 cell line was purchased from ATCC (#HTB-22, ATCC) and was maintained in RPMI-1640 (#11875093, Gibco) with 10% FBS, 1% P/S, and 1% Glutamax (#35050061, Gibco) [30]. The Phoenix-Ampho cell line was purchased from ATCC (#CRL-3213, ATCC) and was maintained in DMEM (#11965092, Gibco) supplemented with 10% FBS and 1% P/S [31]. All cell lines were routinely passaged and maintained in a humidified incubator at 37°C and 5% CO2. Cell line authentication was confirmed using CellCheck 9 Plus (IDEXX BioAnalytics, Westbrook, ME, USA). All cell lines were negative for mycoplasma contamination (MycoAlert Mycoplasma Detection Kit, #LT07-318, Lonza, Basel, Switzerland).

### Human Tissue Collection

The study was approved by the Institutional Review Board (IRB) of Indiana University (Indianapolis, IN, USA; IRB protocol #1812764043). Tissue collection and deidentification were performed by the Biospecimen Collection and Banking Core (BC^2^, Indiana University Melvin and Bren Simon Comprehensive Cancer Center, Indianapolis, IN, USA) with written informed consent. At the time of the hysterectomy, a fresh endometrial specimen was obtained from the uterine fundus away from obvious leiomyomata. The indication for hysterectomy was abnormal uterine bleeding and chronic pelvic pain, thought to be due to previous surgery-proven endometriosis. The patient did not take any medications. At the time of surgery, powder-burn lesions consistent with old endometriosis were biopsied, revealing benign fibrous tissue and smooth muscle on final pathology. The final pathology report of the uterine corpus indicated proliferative-type endometrium and focal adenomyosis. The fallopian tubes and ecto- and endocervix showed no pathological change.

### Primary Culture

Fresh uterine tissue was transferred to the lab in Hanks’ Balanced Salt Solution (HBSS; #14175095, Gibco) with 1% P/S. It was mechanically and enzymatically digested with collagenase/hyaluronidase (#07912, StemCell Technologies, Vancouver, BC, Canada) in DMEM/F12 supplemented with 1% antibiotic-antimycotic (Anti-Anti; #15240062, Gibco). The dissociated cells were passed through a sterile 40-micron filter. The primary cultures were grown in HES.

### Fluorescence-activated Cell Sorting (FACS) of Populations

Cells were tagged with the phycoerythrin (PE)-fluorochrome-conjugated CD326 (EpCam) antibody according to the manufacturer’s protocol specifications. Briefly, cultured cells were trypsinized, collected, and reconstituted to 10^6^ cells per sample in 98 μL sorting buffer (1% charcoal-dextran FBS + 1x PBS + 2mM ethylenediamine tetraacetic acid) and supplemented with 2μl of EpCam-PE antibody (#130-110-999, Miltenyi Biotec, Gaithersburg, MD, USA) or 2 μL of REA isotype control (S)-PE IgG1 antibody (#130-113-438, Miltenyi Biotec). Cells were incubated in a 1:50 antibody/buffer solution x 10min in the dark at 4ᵒC, washed with 1ml of fresh buffer, filtered, and counterstained with propidium iodide (#PI304MP, Thermo Fischer Scientific) at 1:500 concentration in 1ml of the sorting buffer for exclusion of dead cells. Cells were immediately sorted with the BD FACSAria Fusion cell sorter and BD FACSDiva Software (BD Biosciences, Franklin Lakes, NJ, USA) in the Flow Cytometry Resource Facility (FCRF, Indiana University Melvin and Bren Simon Comprehensive Cancer Center, Indianapolis, IN, USA).

### Cell Immortalization

Cells were immortalized via retroviral transduction with the human telomerase reverse transcriptase gene (*TERT*) using a pLXSN-hTERT vector (kindly provided by Dr. Hari Nakshatri, Indiana University [32]). For viral production, 2.5 x 10^6^ Phoenix-Ampho packaging cells were transfected with 20 μg of pLXSN-hTERT DNA with 25 μL of Lipofectamine-2000 reagent (#11668019, Thermo Fisher Scientific) in Opti-MEM (#31985070, Thermo Fisher Scientific) per the manufacturer’s recommendation. After 24 h of incubation at 37°C and 5% CO_2_, the medium was replaced. After 48 h (72h post-transfection), the virus-containing medium supernatant was collected, centrifuged to remove the debris, filtered through a 0.45 μm filter, and kept at −80°C until use. Primary EpCam+ cell cultures were sub-cultured in six-well plates until 50% confluence and infected with viral supernatant in the presence of 4 μg/ml polybrene (#TR-1003-G, Millipore, Burlington, MA, USA). Transformed cells were selected with 300 μg/ml geneticin (#10131027, Thermo Fisher Scientific).

### Authentication and Short-Tandem Repeat Profiling

A cryovial of passage two cells in freezing medium [charcoal-dextran FBS + 10% dimethyl sulfoxide (DMSO; #D8418, Sigma-Aldrich)] was sent to IDEXX BioAnalytics (Columbia, MO 65201, USA) for CellCheck16-Human Test (human 16-marker STR profile and interspecies contamination test).

### Karyotype Analysis

Simple karyotypic analysis was completed for early and late passages using KaryoLogic services (Durham, NC 27713, USA). Cells at early and late passages (p.6 and p.53) were prepared according to their specimen preparation and shipping instructions.

### Immunofluorescence Microscopy

Cells were cultured on Lab-Tek II 8-well chamber slides (#154534PK, Thermo Fisher Scientific), fixed with 4% paraformaldehyde (#J61899-AP, Thermo Fisher Scientific) in PBS for 15 min, rinsed with PBS 3 x 3min, and permeabilized with ice-cold acetone (#A929SK4, Thermo Fisher Scientific) at −20°C x 10 min. Following permeabilization, cells were rinsed with PBS 3 x 3 min at RT, blocked with 5% normal goat serum (#S-10000, Vector Laboratories, Inc., Burlingame, CA) for 1 hour at RT, and incubated with primary antibody **(Suppl. Table S1)** in a blocking solution of 1% BSA (#sc-2323, Santa Cruz Biotechnology, Dallas, TX) with 0.05% Triton X-100 (#T8787, Sigma-Aldrich) in PBS overnight at 4°C. After 3 x 3 min washes with PBS, samples were further incubated with the fluorescent-conjugated secondary antibody at 1:200 dilution, incubated in the blocking solution for 1 h at RT in the dark, and washed again 3 x 3 min with PBS. Slides were further disassembled to remove the wells, DAPI (diamidino-2-phenylindole)-counterstained and mounted using Fluoromount G (#00-4959-52, Invitrogen) overnight. Imaging was conducted using the Advanced Microscopy Group (AMG; Mill Creek, WA) EVOS FL imaging system **(Suppl. Table S1)**.

### Cell Morphologic Analyses and Determination of Appropriate Culture Media

The tAEC21 cells were seeded in triplicate (25,000 cells/well) in 6-well plates and cultured in various growth media **(Suppl. Table S1)** for 7 days with daily fresh growth medium replacement. Daily cell imaging was performed with AMG EVOS FL Imaging System.

### Proliferation Assay

Cells were plated at 500 cells per well of a 96-well plate with each media. Proliferation was assessed using CellTiter 96 AQueous One Solution Cell Proliferation Assay (#G3581, Promega, Madison, WI). Absorbance readings were taken at 490 and 630 nm using a Synergy H1 Hybrid Reader (BioTek, Winooski, VT) and Gen5 3.11 software. Background absorbance was subtracted, and data were normalized to day 1 post-seeding. GraphPad Prism software version 9.2.0 (Dotmatics Platform, Boston, MA, USA) was used for analysis, graphing, and calculation of tAEC21 doubling time in each medium type. All experimental conditions were repeated in biological triplicates. One-way ANOVA with Tukey’s multiple comparison statistical analysis was used to show a significant difference in the proliferation of tAEC21 cells in varying media based on a 95% confidence interval.

### Gene Expression Response to Cytokine Pressure

12Z and tAEC21 cells were seeded at 1.5 x 10^5^ cells/mL in serum-free media overnight. Cells were treated with 15 ng/ml tumor necrosis factor-alpha (TNF-α; #PHC3011, Gibco) or vehicle (DI water) for 24 hours. RNA was isolated using the miRNeasy mini kit (#217004, Qiagen) with an on-column RNase-Free DNase Set (#79254, Qiagen) according to the manufacturer’s protocol. RNA concentration and purity were confirmed with NanoDrop ND-1000 (Thermo Fisher Scientific). Using 1 µg total RNA, complementary DNA (cDNA, High-Capacity cDNA Reverse Transcription Master Mix with RNase Inhibitor, Thermo Fischer Scientific) was created. Samples were diluted to 100 μL and 2 μL was used for qPCR. qPCR was performed using SYBR Green Universal PCR Master Mix (#4309155, Thermo Fisher Scientific) and gene-specific primers **(Suppl Table S1)**. Reactions were run on a QuantStudio-3 Real-Time PCR System (Thermo Fisher Scientific). The primer oligonucleotide pairs were obtained from Millipore Sigma (Burlington, MA, USA). The relative quantification (RQ) of mRNA expression was calculated according to the 2^-ΔΔCt^ method [33]. All experiments were conducted in technical and biological triplicates; data were analyzed with a Mann-Whitney U-test or unpaired Student’s t-test as specified in figure legends (statistical significance defined as p<0.05) using GraphPad Prizm version 9.2.0.

### 3D Spheroid Assembly

For monotypic spheroid culture, tAEC21 cells were seeded at 14,000 cells/well in a U-bottom ultra-low adhesion 96-well plate (S-bio, MS-9096 UZ, Hudson, NH) and centrifuged at 2000 rpm x 3 min. For heterotypic spheroid co-culture, tAEC21 and THESC cells were mixed in a 1:1 cell ratio, seeded in U-bottom plates, and centrifuged. 3D cultures were incubated at 37°C, and spheroid assembly was imaged at 24 h using an AMG EVOS imaging system. The spheroid diameter was determined using ImageJ software (NIH free download) [34, 35]. Statistical analysis was performed using a two-sample unpaired Welch’s t-test (statistical significance defined as p<0.05; n≥39) in GraphPad Prizm software version 9.2.0.

### Transient Fluorescent Staining and 3D Co-Culture Spheroid Assessment

Green 5-chloromethylfluorescein diacetate (CMFDA; #C7025, Invitrogen by Thermo Fisher Scientific) and red 4-([4-(chloromethyl)phenyl]carbonylamino)-2-(1,2,2,4,8,10,10,11-octamethyl-10,11-dihydro-2H-pyrano[3,2-g:5,6-g′]diquinolin-1-ium-6-yl)benzoate (CMTPX; #C34552, Invitrogen by Thermo Fisher Scientific) CellTrackers were utilized for short-term fluorescent labeling of tAEC21 and THESC cells, respectively, as published previously [36]. Briefly, each CellTracker was diluted with dimethyl sulfoxide (DMSO; #D8418, Sigma-Aldrich, Saint Louis, MO, USA) to a 2 mM stock concentration. tAEC21 and THESC cell monolayers were grown to 70% confluence, washed with serum-free medium (SFM), and incubated in SFM with green CMFDA (1:500 ratio; 4 µM final dye concentration) or red CMTPX (1:1000 ratio; 2 µM final dye concentration), respectively. After 30 min at 37°C, the CellTracker was removed from the cells; cells were incubated for an additional 30 min at 37°C in complete media; fluorescence was confirmed using the AMG EVOS FL imaging system. To generate 3D heterotypic spheroids, green tAEC21, and red THESC cells were trypsinized, mixed in a 1:1 cell ratio, seeded (14,000 cells/well) in a U-bottom ultra-low adhesion 96-well plate (S-bio, MS-9096 UZ), and centrifuged at 2000 rpm x 3 min. Co-cultures were incubated at 37°C and imaged at 0 h (immediately after centrifugation), 24, 48, and 72 h using AMG EVOS FL microscope.

### Spheroid Fixation and Histological Analysis

Spheroids (three days post-seeding) were collected, briefly rinsed with 1x PBS, fixed in 4% paraformaldehyde (Thermo Fisher Scientific) x 30 min, re-rinsed with 1x PBS, and stored in 70% ethanol for an additional 30 min or until embedding. For embedding, spheroids were placed in 1.5 ml Eppendorf tubes and covered with 300 µl of pre-warmed Epredia HistoGel specimen processing gel (#22110678, Thermo Fisher Scientific) and allowed to solidify for 20 min on ice. The HistoGel plug with trapped spheroids was then gently removed from the tube with a spatula, immediately placed in a tissue embedding cassette in 70% ethanol, and transferred to the Histology Core of the Indiana Center for Musculoskeletal Health (Indiana University School of Medicine, Indianapolis, IN USA) for processing, paraffin embedding, 5 µm sectioning, and hematoxylin and eosin (H&E) staining. Additional unstained sections were sent for immunostaining.

### Immunohistochemistry (IHC)

Spheroid tissue immunostaining was performed by the Immunohistochemistry Research Core (Indiana University School of Medicine, Indianapolis, IN USA) on an Agilent Dako Omnis platform (Santa Clara, CA, USA). Briefly, sections on slides underwent deparaffinization, rehydration, heat-induced epitope retrieval (Agilent PT Link), antigen retrieval (Dako EnVision FLEX Target Retrieval Solution, high pH), followed by quenching with 3% H_2_O_2_ x 3 min for endogenous peroxidase (HRP) activity elimination. Primary antibody incubation, optional signal amplification (Dako Envision FLEX + Mouse or Rabbit LINKER), detection (Dako Envision FLEX/HRP), 3,3’-diaminobenzidine visualization (Dako EnVision FLEX DAB+ Chromogen), and counterstaining (Dako EnVision FLEX Hematoxylin) were performed as specified in **Suppl.Table S1**, followed by dehydration, clearing, and coverslip mounting. Imaging was performed using a Zeiss Axio Lab.A1 Microscope and Labscope software (Carl Zeiss Microscopy LLC; Oberkochen, Germany; **Suppl. Table S1**).

## RESULTS

### Adenomyosis epithelial cell line establishment, authentication, and genetic evaluation

After mechanical and enzymatic dissociation of the fresh endometrial tissue, cultured primary cells displayed morphological heterogeneity with secluded islets of epithelial-like polygonal cells surrounded by fibroblast-like stroma **(Figure 1A**, left panel). Cell sorting using epithelial cell adhesion molecule (EpCAM), a cell surface marker for epithelial cells, allowed successful separation of epithelial (EpCAM+) and non-epithelial (EpCAM-) cell subpopulations **(Figure 1A**, center and right panels; **Suppl. Figure 1A)** [37]. Epithelial-like EpCAM+ cells underwent retroviral transduction of the human telomerase (hTERT) gene, selection with geneticin, and cell expansion. The cell line was designated tAEC21 (TERT-immortalized adenomyosis epithelial cells from patient 21) cell line. Via qPCR, hTERT expression in the tAEC21 cells was as high as THESC (365±47-fold expression increase) **(Suppl. Figure 1B)**. To date, the tAEC21 cell line has been successfully passaged >80 times with no proliferation decline or loss of epithelial-like morphology in vitro (data not shown).

**Figure 1.**
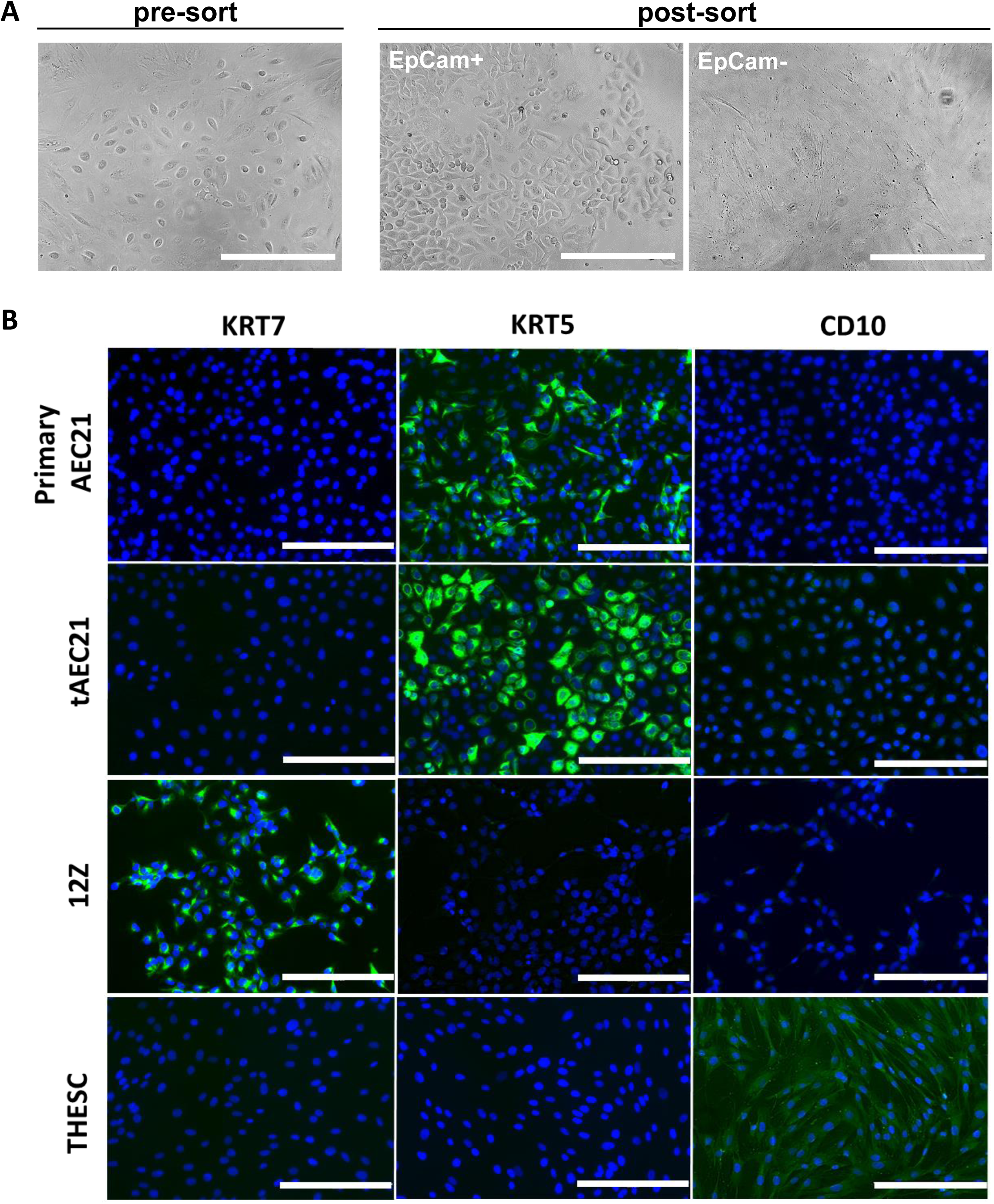
A newly established tAEC21 cell line possesses morphology and marker expression consistent with epithelial-like phenotype. (A) A heterogeneous primary cell population dissociated from fresh endometrial tissue of an adenomyosis patient was cultured in vitro. After sorting, EpCam+ (epithelial) and EpCam-cell subpopulations. Micrographs: AMG EVOS fluorescence microscope; scalebar: 200μm. (B) Primary AEC21, hTERT-immortalized tAEC21, epithelial-like 12Z, and stromal fibroblast THESC cells were cultured on 8-well chamber slides and evaluated for cytokeratin-7 (KRT7), cytokeratin-5 (KRT5), and CD10 immunofluorescence, nuclei-counterstained with DAPI (blue), and imaged using an AMG EVOS fluorescence microscope; scalebar: 200μm.

Morphologically in vitro, the tAEC21 cells appeared very epithelial-like (**Figure 1A**, center panel). To further examine the basic molecular epithelial features, we performed immunofluorescent staining of primary cultures and the immortalized tAEC21 line. We used published data from the endometriotic epithelial-like 12Z line to guide our cytokeratin(KRT)-7 marker selection [38]. Initial immunofluorescent staining of primary cells and the immortalized tAEC21 line showed no KRT7 expression (**Figure 1B**, left panel). Subsequent panel expansion to address other commonly explored endometrial epithelial cytokeratins KRT5, KRT8, KRT19, KRT20 (reviewed in [39]) detected the expression of a basal epithelial marker KRT5 in both primary and tAEC21 cells (**Figure 1B**, center panel). As the most likely contaminating cell type would be stromal cells, we examined the stromal cell marker CD10 using published data from the stromal cell line, THESC [38, 40]. Both primary cells and the immortalized tAEC21 line were CD10-negative (**Figure 1B**, right panel). The tAEC21 cell line exhibits properties of epithelial cells and not stromal cells.

To further characterize the tAEC21 cell line, we examined additional markers. Again, we used published data from the endometriotic epithelial-like cell line to guide our marker selection [22]. In human adenomyosis, neuronal cadherin (Ncad) is expressed in both eutopic and ectopic endometrial tissues at higher levels compared to normal endometrium [41, 42]. On the other hand, the decrease or loss of epithelial cadherin (Ecad) is observed in both eutopic and ectopic adenomyosis uterine tissues in parallel with increased expression of Ncad [42–44]. This cadherin switch is associated with the epithelial-to-mesenchymal transition (EMT) and is typically accompanied by upregulated expression of vimentin, a canonical marker of mesenchymal lineages or EMT-reprogrammed epithelial cells with enhanced invasive properties [45]. Assessed via immunostaining, both primary culture and immortalized tAEC21 cells were strongly positive for junctional Ncad (**Figure 2**). This Ncad staining was consistent with the staining in 12Z cells (**Figure 2**), as previously published [25, 46]. Primary and immortalized tAEC21 cells were negative for Ecad (**Figure 2**). Similarly to previously reported, 12Z cells were also negative (**Figure 2**) [25, 46]. Further, primary cultures exhibited strong cytoplasmic (predominantly perinuclear) vimentin expression in roughly 50% of the cells and moderate vimentin staining in the other half of the cells (**Figure 2**). The tAEC21 cells exhibited less frequent, highly positive cytoplasmic vimentin staining, with most cells displaying weak staining. 12Z cells exhibited a small portion of strongly positive cells and the rest of cells were weakly positive for perinuclear vimentin (**Figure 2**). A non-invasive breast cancer MCF-7 cell line with poor metastatic potential served as an E-cadherin-positive, N-cadherin- and vimentin-negative antibody control (**Figure 2**) [47].

**Figure 2.**
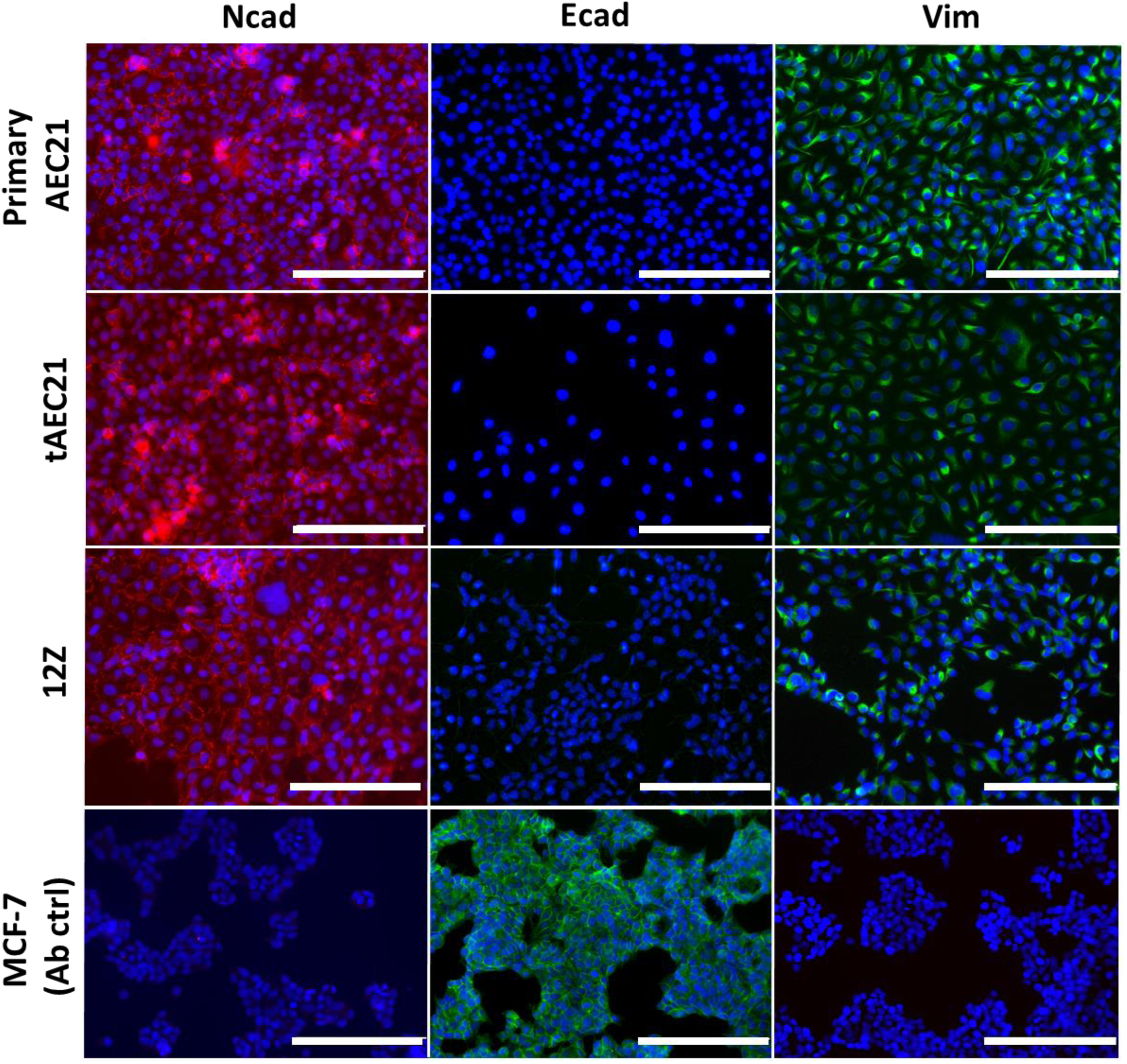
The tAEC21 cell line exhibits epithelial-mesenchymal transition markers. Primary AEC21, hTERT-immortalized tAEC21, epithelial-like 12Z, and MCF-7 (antibody ctrl) cells were cultured on 8-well chamber slides and evaluated for N-cadherin (Ncad), E-cadherin (Ecad), and vimentin (Vim) immunofluorescence, nuclei-counterstained with DAPI (blue), and imaged using an AMG EVOS fluorescence microscope; scalebar: 200μm.

Short-tandem repeat profiling is a rigorous method of authenticating a new cell line [48]. We performed CellCheck 16-Human Test (IDEXX) on the tAEC21 cell line. This commercially available service showed that the tAEC21 is a human-derived cell line, without interspecies contamination. IDEXX provided the 16-marker short tandem repeat (STR) profile for tAEC21 (**Table 1**) with a comparison to the DSMZ STR database [49]. The STR profile of tAEC21 did not match any lines in the database and was not a cross-contaminant or misidentified cell line.

**Table 1.** IDEXX CellCheck^TM^ 16 established a unique 16 short-tandem repeat (STR) marker profile for tAEC21 cell line.

| Marker Name | tAEC21 Genetic Profile Matches |
| --- | --- |
| *AMEL* | X |
| *CSF1PO* | 11 |
| *D13S317* | 8, 10 |
| *D16S539* | 11, 12 |
| *D18S51* | 16, 18 |
| *D21S11* | 31, 31.2 |
| *D3S1358* | 14 |
| *D5S818* | 11, 12 |
| *D7S820* | 11, 12, 13, 14 |
| *D8S1179* | 14, 15 |
| *FGA* | 23, 24, 25, 26 |
| *Penta_D* | 12, 13 |
| *Penta_E* | 5, 13 |
| *TH01* | 9, 9.3 |
| *TPOX* | 8, 11 |
| *vWA* | 18, 19 |

### Therefore, tAEC21 is a novel, immortalized cell line

Typical karyotyping involves creating a metaphase spread of the chromosomes, aligning each chromosome pair by its characteristic banding pattern, and counting the number of chromosomes in metaphase. A normal human cell would be expected to have 23 pairs or 46 chromosomes [50, 51]. To do this in tAEC21 cells, we prepared and shipped tAEC21 cells at passages 6 (early) and 53 (late), to KaryoLogic. KaryoLogic could only get 10 metaphase spreads each from the early and late passages. Typically, 20 metaphase spreads are examined for a typical karyotype. Data output from KaryoLogic showed that multiple chromosomal rearrangements were observed, making the alignment of pairs impossible. Further, all 10 of the metaphase spreads from early and late passages showed more than 46 chromosomes. The 10-metaphase chromosome counts at early passage were (76, 77, 79, 81, 81, 81, 82, 82, 83, and >100) and at late passage were (67, 69, 71, 72, 73, 74, 75, 76, 77, and >100) passages. All of the cells were polyploid. Representative metaphase spread images are displayed (**Supplementary Figure S2**).

### Optimized Cell Growth Media

The development of a novel cell line requires the establishment of cell growth patterns and the optimization of culture conditions that are suitable for diverse experimental applications [52]. The tAEC21 cells were maintained in the most commonly used, commercially-available basal cell culture media (**Suppl. Table S1**) supplemented with 10% FBS and 1% P/S. Cellular morphology and confluency were monitored daily. Microscopically, the tAEC21 cells demonstrated consistent and comparable growth in DMEM/F12 and RPMI-based complete media. In both media types, cells avidly propagated, forming a fully confluent monolayer by day 7 (168 h), while retaining regular epithelial morphology (**Figure 3A**, left and middle panels). In contrast, tAEC21 cells cultured in the MEM-based media lost their characteristic polygonal morphology, acquired spindle-like cell shape, and proliferated more slowly (**Figure 3A**, right panel). To examine the effects of specific serum derivations, tAEC21 cells were grown in DMEM/F12 basal medium supplemented with charcoal-dextran-stripped fetal bovine serum (c/d FBS). DMEM/F12 c/d FBS-cultured tAEC21 cells retained their original epithelial morphology and exhibited a steady growth rate, reaching confluency by day 7 (**Suppl. Figure S3A**).

**Figure 3.**
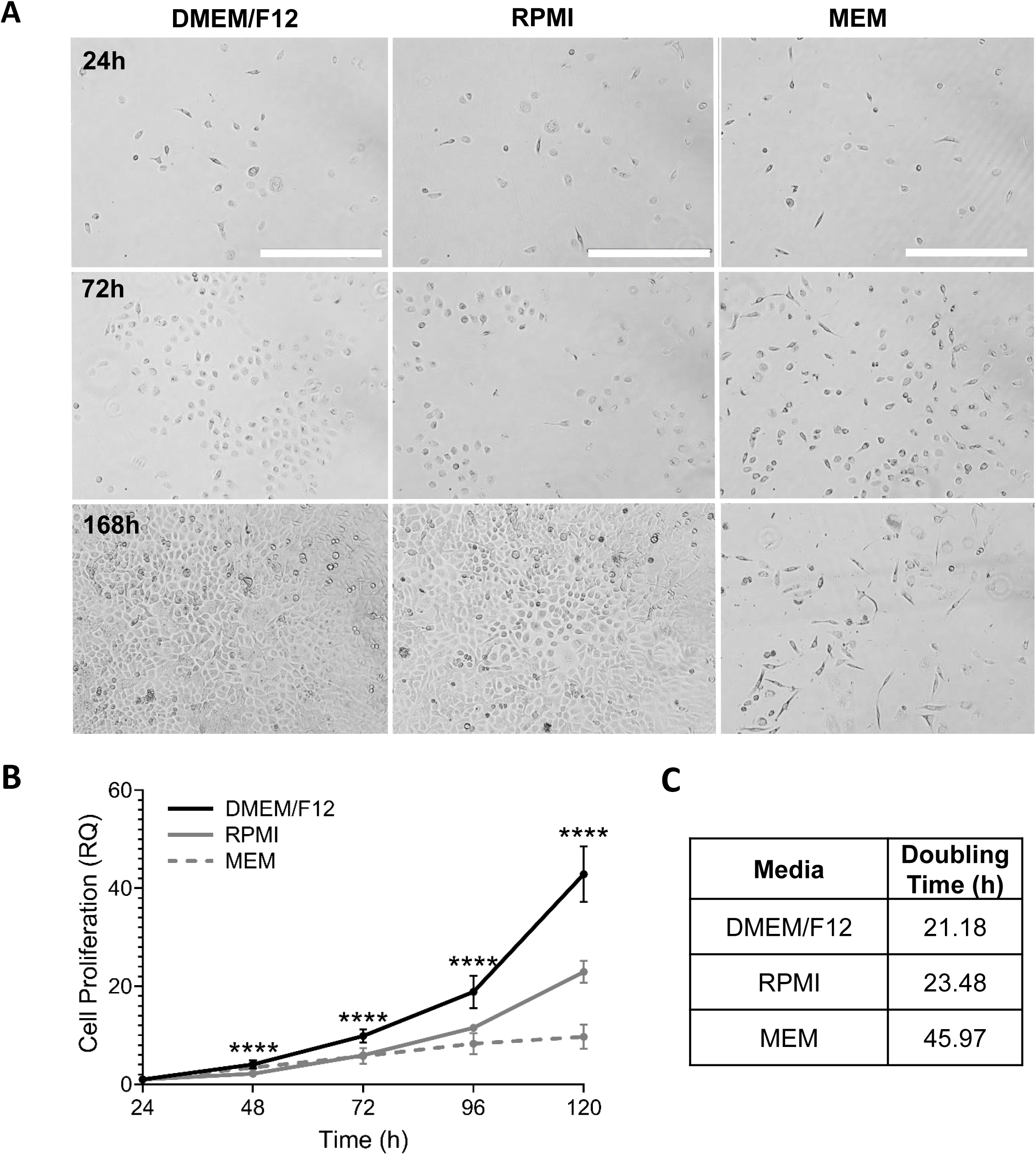
The tAEC21 cell line grows in common commercially available media. A) tAEC21 cells were seeded (25,000 cells/well) in a 6-well plate in DMEM/F12, MEM, or RPMI supplemented with 10% FBS and 1% P/S. Cell growth/morphology was monitored for 168 h (7 days) with daily fresh medium exchange (n=3). Images were taken using AMG EVOS FL imaging system; scale bar: 400 μm. B) tAEC21 were plated (500 cells/well) in a 96-well plate and subjected to MTS proliferation assay. Data are presented as relative quantification (RQ) with absorbance values normalized to 24 h post-seeding. Statistical analysis (one-way ANOVA with Tukey’s multiple comparison; ****, p < 0.0001; n=3) and C) cell doubling time acquisition (nonlinear fit) were performed in GraphPad Prism version 9.2.0.

To more objectively measure cellular proliferation, the CellTiter 96 AQueous One Solution Cell Proliferation Assay (#G3581, Promega, Madison, WI) was used. The highest tAEC21 proliferation rate was found in DMEM/F12-based medium compared to other media types (p<0.0001, Turkey multiple comparison test) (**Figure 3B; Suppl. Figure S3**). DMEM/F12 showed the fastest doubling time at 21.2h, while MEM-based media was nearly double that doubling time at 46h (**Figure 3C; Suppl. Figure S2C**).

### Hormone receptor status and endometrial molecule expression

Adenomyosis is a hormone-dependent disease associated with increased estrogen receptor expression (predominantly estrogen receptor 1, ESR1; also known as ERα) and simultaneous progesterone resistance [53–55]. Assessing the estrogen and progesterone receptor (PGR) status in a newly established cell line is essential for evaluating its hormonal sensitivity and potential utility in disease modeling. The tAEC21 cells cultured in monolayer showed no ERα or PGR expression, and findings mirrored those in endometriotic epithelial-like 12Z cell monolayers (**Figure 4A**). As previously published, MCF-7 breast cancer cells stained strongly positive for both receptors (**Figure 4A**) [47]. To expand tAEC21 cell line molecular characterization outside the endometrium, we evaluated the expression of paired box gene 8 (PAX8), a transcriptional factor implicated in embryonic development of Müllerian tissues and known to be expressed in both normal eutopic and ectopic (adenomyotic/endometriotic) epithelial endometrium, as well as Müllerian neoplasms [56–58]. Separately, the status of AT-rich interactive domain-containing protein 1A (ARID1A), a chromatin remodeler and a tumor suppressor often mutated in deep infiltrating endometriosis and endometriosis-associated malignancies, was assessed. Both tAEC21 and 12Z cell lines displayed high nuclear expression of PAX8 and ARID1A proteins (**Figure 4B**), with endometrial stromal THESC and ovarian clear cell carcinoma TOV21G cells utilized as respective PAX8-and ARID1A-negative controls [56, 59, 60].

**Figure 4.**
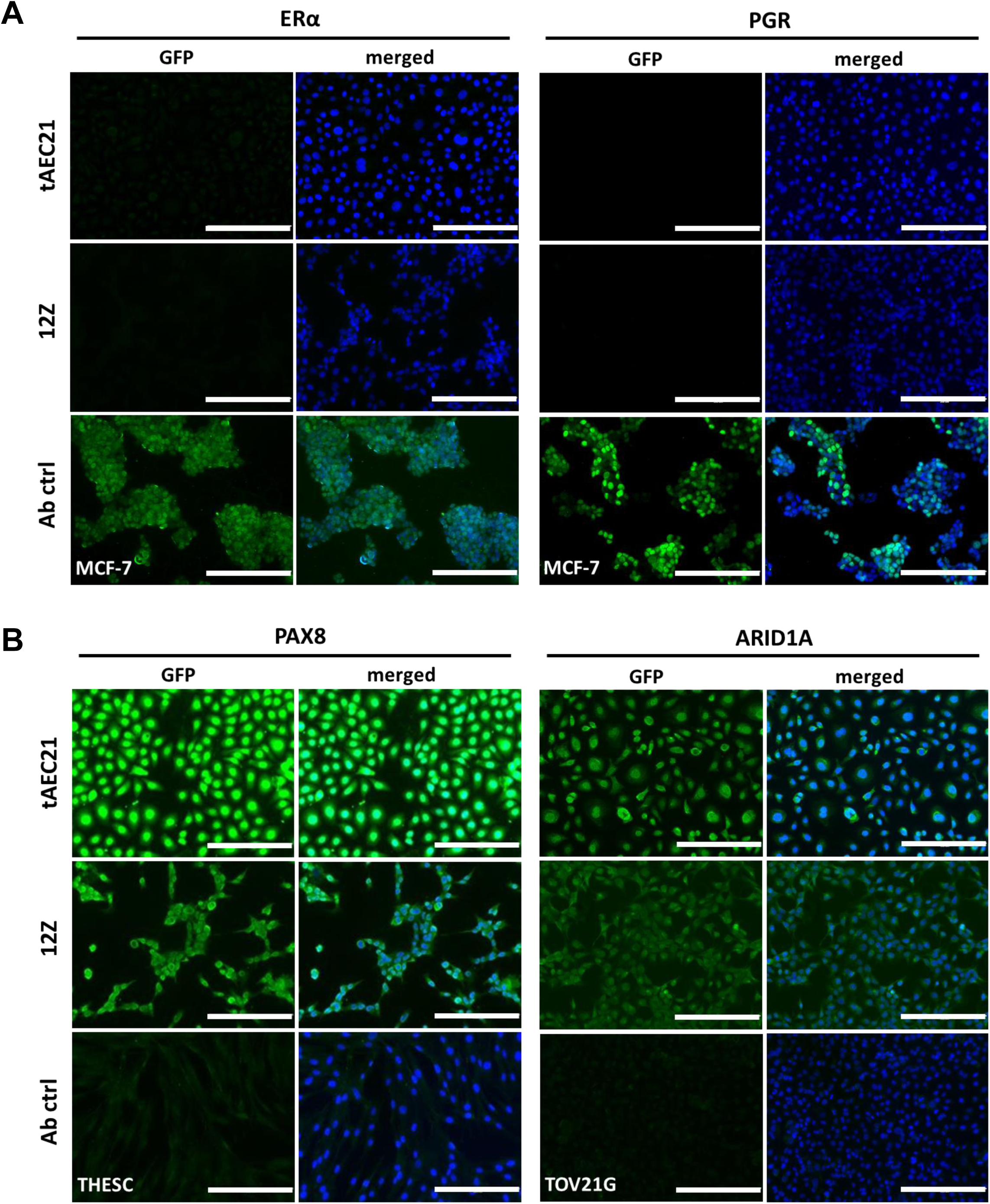
The tAEC21 cell line is negative for estrogen and progesterone receptors. tAEC21, 12Z, MCF-7 (positive ERα/PGR antibody control), THESC (negative PAX8 antibody control), TOV21G (negative ARID1A antibody control) cells were subcultured on 8-well chamber slides and processed for (A) estrogen (ESR1/ERα) and progesterone (PGR) receptors, (B) paired box (PAX)8, and AT-rich interaction domain 1A (ARID1A) immunofluorescence, nuclei-counterstained with DAPI (blue), and imaged using an AMG EVOS fluorescence microscope; scalebar: 200μm.

### Response to cytokine stimulation

Abnormal levels of pro-inflammatory mediators, such as tumor necrosis factor (TNF)-α, interleukin (IL)6, C-X-C motif chemokine ligand 8 (CXCL8; also known as IL8), and C-C motif chemokine ligand 2 (CCL2; also known as monocyte-chemoattractant protein 1, MCP-1), are consistently elevated in the adenomyotic uterus, and peritoneal fluids of women with adenomyosis [54, 61, 62]. To determine the initial suitability of a newly created cell line for modeling disease-associated inflammation, we evaluated the tAEC21 line’s sensitivity to TNF-α stimulation. TNF-α-treated tAEC21 cell monolayers showed notably enhanced *IL6* (8.8±1.4-fold increase), *CXCL8* (11.8±2.1-fold increase), and *CCL2* (14.4±1.6-fold increase) gene expression (mean ± SEM, p<0.0001) (**Figure 5**). A moderate but statistically significant upregulation of mucin 1 (*MUC1*), a TNF-α-induced transmembrane mucin constitutively expressed in normal uterine and ectopic endometrium, was observed (3.0±0.2-fold increase, p<0.0001) (**Figure 5**) [63, 64]. The altered expression pattern was consistent with a previously reported TNF-α-sensitive 12Z cell line, which was a TNF-α response control in current experiments (**Figure 5**) [38, 65].

**Figure 5.**
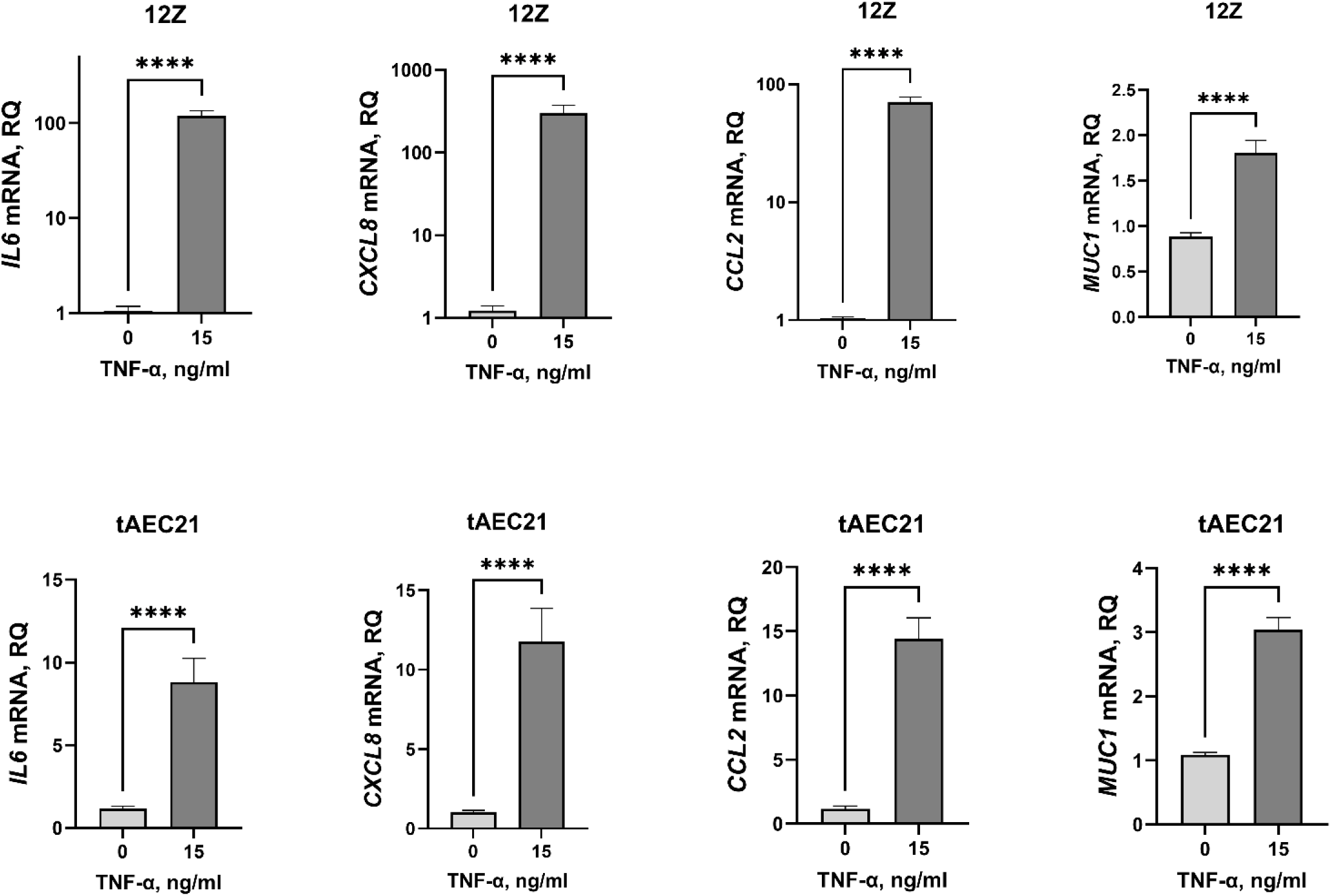
The tAEC21 cell line elicits an immune response to TNF-α cytokine pressure. tAEC21 and 12Z cell monolayers were treated with TNF-α (15 ng/ml) or vehicle (water) for 24h, and RNA was extracted and processed for qPCR. Data are presented as relative quantification (RQ) of indicated gene transcript expression in TNF-α-treated cells compared to vehicle-treated cells, normalized to β-actin endogenous control (mean ± SEM, n≥3); ****, p<0.0001, Mann-Whitney U-test. Statistical analysis was performed in GraphPad Prism (version 9.2.0). *IL6* – interleukin 6; *CXCL8* –C-X-C motif chemokine ligand 8; *CCL2* – C-C motif chemokine ligand 2; *MUC1* – mucin 1.

### Three-dimensional (3D) spheroid architecture and molecular profile

Two-dimensional (2D) monolayer cell cultures have been foundational for *in vitro* disease modeling, offering simplicity and high-throughput screening capabilities. However, 3D culture approaches are more relevant for recapitulating the complex multicell-type tissue environment, providing a more accurate representation of cell-cell and cell-matrix interactions [66]. To explore their 3D dynamics and physiological relevance, tAEC21 cells were grown in 3D. They were grown alone as monotypic epithelial spheroids and with THESC as heterotypic spheroids. In 3D, tAEC21 cells formed larger (>1500 μm in diameter), loosely attached multicellular aggregates (**Figure 6A**, left and right panels). Co-culture of tAEC21 with THESC endometrial stromal fibroblasts resulted in spatially organized, cohesive spheroid structures of significantly smaller diameter (roughly 500 μm in diameter) (**Figure 6A**, center and right panels).

**Figure 6.**
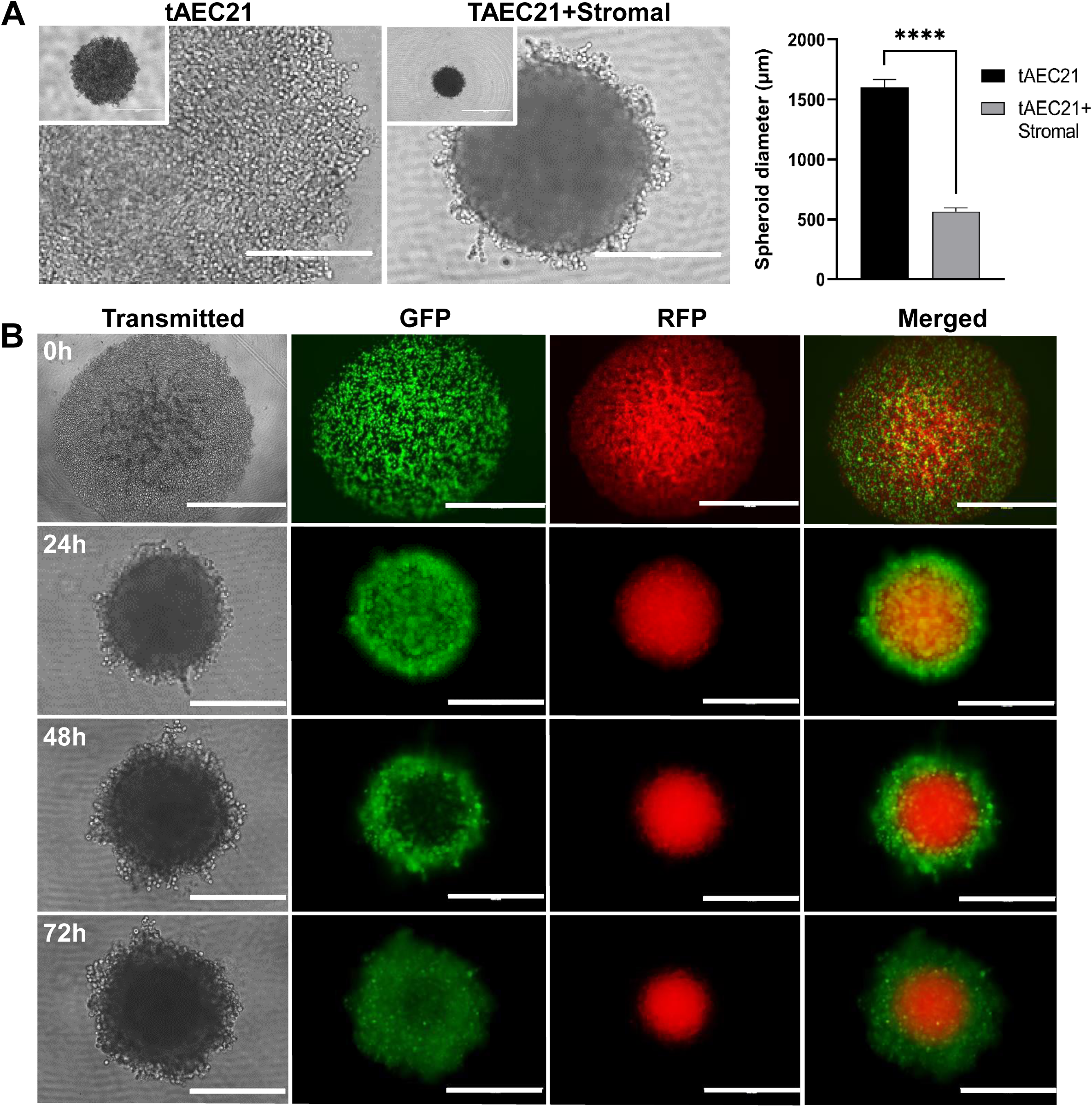
Epithelial-like tAEC21 cells and supporting stroma assemble into a biologically relevant pattern in a 3D co-culture model. A) tAEC21 were seeded into spheroids alone (14,000 cells/well) or in co-culture with stromal THESC cells (14,000 cells/well; 1:1 cell ratio) and imaged with the EVOS FL imaging system at 24h; scalebar: 1000 μm. B) CMTPX-tagged (red) stromal THESC and CMFDA-tagged (green) tAEC21 cells were co-cultured (14,000 cell/well, 1:1 cell ratio) in a U-bottom 96-well plate and spheroid conglomeration was monitored with the EVOS FL imaging system at 0-72h in transmitted, green fluorescent protein (GFP) and red fluorescent protein (RFP) modes. Experiments were performed in triplicate and representative images are shown; scalebar: 1000 μm. ****p <0.0001 (two-sample unpaired Welch’s t-test).

Immunofluorescent imaging of live heterotypic 3D spheroids composed of transient fluorophore-tagged epithelial-like tAEC21 (green, CellTracker) and stromal, THESC (red, CellTracker) over time showed a physiologic response of cells. Over time, the epithelial cells formed a shell around the stromal core. At 0 hours, the green epithelial cells were intermingled with the red stromal cells. By 24 hours, a green epithelial shell was forming around the red stromal core. Within 48-72h, the green epithelial shell and the red stromal core were well-defined (**Figure 6B**).

To evaluate further, monotypic tAEC21 spheroids and heterotypic tEAC21 with THESC spheroids were fixed, embedded, and underwent molecular immunohistochemistry. During histological processing, loose monotypic tAEC21 multicellular aggregates failed to maintain structural integrity and were disassembled into unorganized clusters (**Figure 7A-B**). Heterotypic spheroids maintained structural integrity and were tightly compacted. Hematoxylin and eosin-stained sections from heterotypic spheroids showed that the epithelial tAEC21 component (evident by round morphology, lightly eosinophilic cytoplasm, and basophilic nuclei) populated predominantly along the periphery around the stromal (spindle-shaped, eosinophilic cells with elongated nuclei) THESC supportive background. Immunostaining confirmed cytokeratin-positive and CD10-negative tAEC21 localization in a spheroid shell (**Figure 7B**). Some epithelial cells interspersed among stroma exhibiting incomplete cell type segregation (**Figure 7A-B**). In alignment with endometrial epithelium, as monotypic spheroids, tAEC21 cells were strongly positive in nearly all cells for hepatocyte nuclear factor 1 beta (NHF1β) and constituted an NHF1β-positive outer shell in the heterotypic co-cultures (**Figure 7B**) [67]. Further, tAEC21 immunostaining for napsin A, a marker for endometrial-associated malignancies, detected no expression (**Figure 7B**) [68]. Importantly, in both monotypic and heterotypic 3D tissue structures, tAEC21 cells expressed ERα but not PGR (**Figure 7B**).

**Figure 7.**
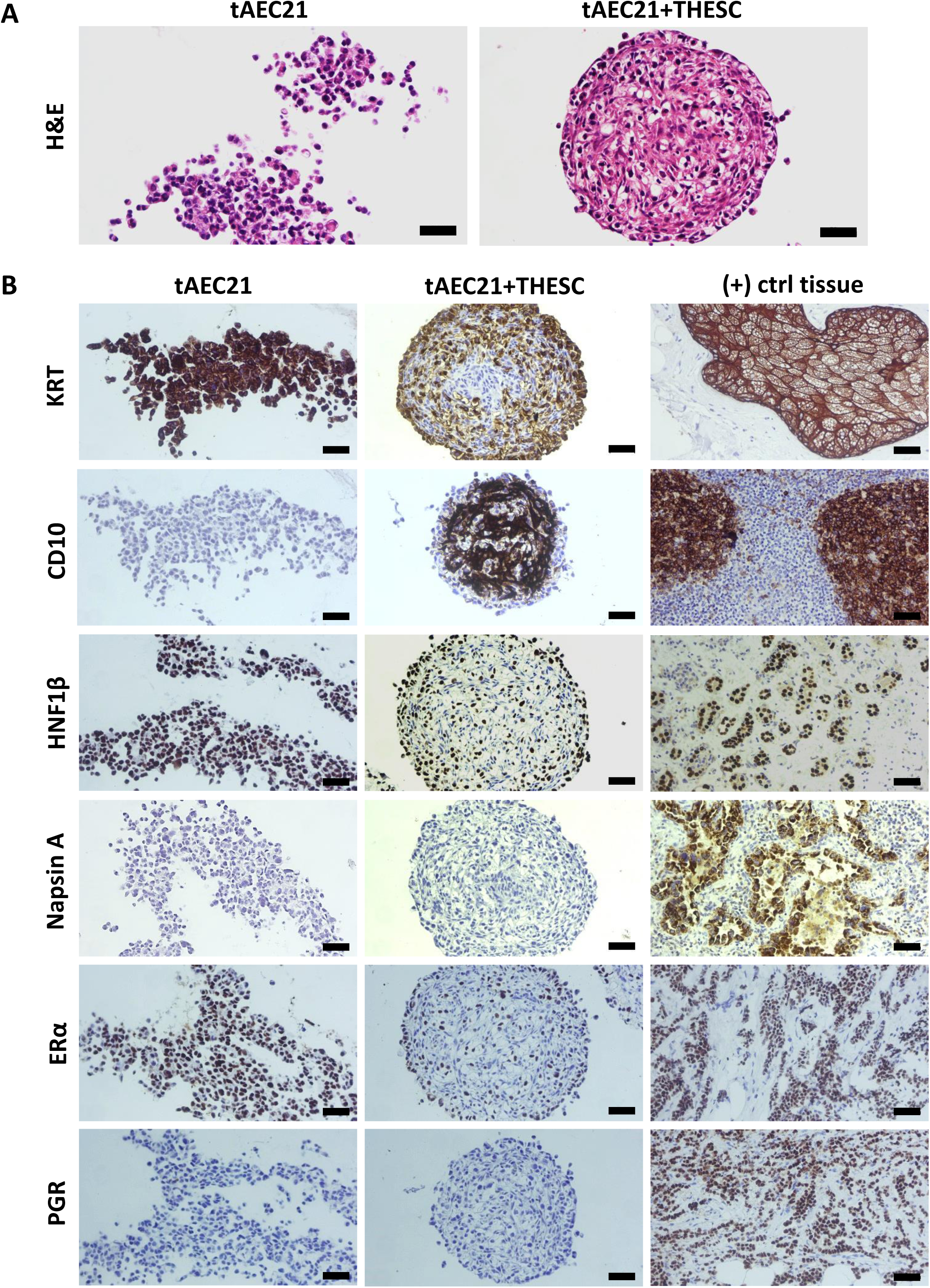
tAEC21/stromal 3D co-cultures exhibit histology and molecular expression patterns consistent with endometrial-like tissue. A) tAEC21 were seeded into spheroids alone or in a mixture with stromal THESC cells (1:1 cell ratio); assembled 3D spheroids were harvested at 72h, fixed, paraffin-embedded, sectioned, and stained with H&E. Scalebar: 50 μm. (B) 3D spheroid tissue sections were subjected to immunohistochemistry and visualized with 3,3’-diaminobenzidine (brown, positive) using a Zeiss Axio Lab.A1 Microscope. Positive (+) control tissues: skin (keratin; KRT+), tonsil (CD10+), renal cell carcinoma (hepatocyte nuclear factor 1 homeobox B; HNF1β+), lung adenocarcinoma (Napsin A+), breast (estrogen receptor; ERα+), breast (progesterone receptor; PGR+). Scalebar: 50 μm.

## DISCUSSION

Etiopathogenesis of adenomyosis remains obscure. Leading proposed theories of the disease development include: a) repetitive microtraumas and disruption of the junctional zone (JZ) between endometrium and subendometrial myometrium followed by invagination of basal endometrial cells into the muscular layer with their subsequent migration and proliferation; b) development of intramyometrial endometrial-like lesions from displaced Müllerian system embryonic pluripotent residues; c) aberrant migration and differentiation of adult endometrial epithelial and stromal progenitor cells, relocated from the endometrium basalis stem cell niches through the disrupted JZ; d) “outside-to-inside” theory, wherein transcoelomic endothelial cells, mislaid after retrograde menstruation or migrating from ectopic endometriosis lesions, penetrate inward through uterine serosa into the myometrium to establish endometrial-like implants [44, 69, 70]. Intramyometrial adenomyotic lesion formation by one or more of the proposed mechanisms occurs amid and is exacerbated by a dysregulated hormone background (imbalance between elevated estrogen levels and loss of progesterone receptivity) [54, 55]. Increased estrogen exposure and persistent tissue injury and repair promote chronic local and systemic inflammatory response and neoangiogenesis, facilitating disease progression [70–72].

Importantly, adenomyosis shares overlapping pathophysiology with endometriosis, as both conditions are characterized by ectopic growth of histologically similar endometrial-like lesions arising from the same origin (uterine endometrium), harbor common molecular abnormalities (in particular, KRAS mutations and various epigenetic and immune alterations), possess estrogen dependence and progesterone resistance, share clinical presentation and are often concurrent [73–75]. Multiple next-generation sequencing studies have identified recurring KRAS mutations in adenomyosis epithelial cells and adjacent basalis endometrial glands, supporting the invagination theory of pathogenesis [76]. These mutations are also observed in endometriosis, suggesting a similar disease process in both conditions [76].

Our newly created hTERT-immortalized adenomyosis-derived epithelial cell line tAEC21 exhibits features of epithelial-like cells, including polygonal morphology, tight packing during monolayer growth, and cell surface expression of epithelial marker EpCam (**Figure 1A; Suppl. Figure S1A**) [37]. The cytoplasmic expression of cytokeratin and the absence of CD10 confirm tAEC21 epithelial-like status (**Figure 1B**) [39, 40]. The cells also stain strongly positive for nuclear PAX8 and nuclear HNF1β (both are markers exclusive to endometrial glandular epithelium, not stroma), confirming their epithelial origin within the endometrium (**Figure 4B**; **Figure 7B**) [56–58, 67].

In addition to epithelial characteristics, tAEC21 cell line possesses hallmarks of EMT (complete loss of Ecad, strong junctional Ncad expression, and strong-to-moderate cytoplasmic vimentin expression) (**Figure 2**). In human normal uterine proliferative-phase endometrium, Ncad is present in 11-20% of epithelial cells and highlights a subset of cells with high self-renewal capacity and proliferative potential, serving endometrial regeneration [43, 77]. Ncad-positive cells are predominantly localized in endometrium basalis, adjacent to the myometrium, and can differentiate into gland-like structures in vitro [77]. Ncad expression has also been linked to increased migration and invasiveness in endometrial-related pathologies through the mechanisms of EMT [22, 39]. On the other hand, Ecad is a key epithelial cell marker that maintains the non-invasive epithelial cell phenotype, epithelial cell-cell adhesion, and layer integrity (reviewed in [78]). Loss of Ecad expression contributes to the EMT shift and the acquisition of a more invasive, mesenchymal cell phenotype [78–80]. Immunohistochemical studies demonstrate clear membranous Ecad staining in human normal endometrial epithelium, with strong expression consistent across the proliferative and secretory phases of the menstrual cycle [41, 43]. Immunostaining data on vimentin expression in human normal endometrial epithelium vary. Dabbs et al. reported perinuclear vimentin staining in normal human proliferative endometrial glands and vimentin absence in secretory phase [81]. Nonwitz et al. showed consistent co-expression of vimentin and cytokeratin in surface and glandular epithelial cells across all menstrual cycle phases [82]. Cyclic expression of epithelial vimentin (predominantly in the proliferative phase with highly variable staining intensity) was also reported in both eutopic and ectopic endometrial tissues of women with adenomyosis [83]. On the contrary, immunostaining performed by Zhou et al. showed low epithelial vimentin in normal uterine endometrial glands and its increased expression in eutopic endometrium and more significantly, ectopic lesions from adenomyosis patients [42]. In alignment with human data, EMT events are observed in murine models of adenomyosis [84]. In genetically engineered mice with constitutive activation of uterine β-catenin, epithelial cells of developed adenomyosis lesions showed Ecad suppression together with expression of vimentin and other EMT-related markers compared to control mice uteri [15]. Similarly, decreased Ecad and increased vimentin immunostaining were observed in ectopic adenomyotic tissues of tamoxifen-induced adenomyosis mice [85].

The described molecular switch is observed in the tAEC21 cells and suggests acquisition of EMT-related cell properties by the cell line, such as increased motility, invasiveness, and altered adhesion [25, 36, 43, 45, 80, 86]. This invasive behavior may explain why epithelial-origin primary cells were initially able to attach to the tissue culture plastic with the pool of stromal cells and were ultimately isolated. Typically, endometrial epithelial cells require extracellular matrix coatings, such as Matrigel, on culture dishes to attach and grow effectively, as this mimics their dependence on the basement membrane for in vivo adhesion [87, 88].

Epithelial-like cells that have transitioned through EMT often gain enhanced cell-substrate adhesion and invasiveness similar to mesenchymal cells and metastatic carcinoma cells, allowing facilitated attachment to more rigid surfaces like uncoated tissue culture plastic [25].

An unexpected finding in tAEC21 cell line characterization is its abnormal polyploid karyotype with multiple chromosomal rearrangements (**Suppl. Figure S2**). Limited studies involving the one published adenomyosis-derived cell line and patient-derived organoids (from 3 different adenomyosis patients) report a normal 46, XX karyotype [27, 89]. Nevertheless, chromosomal alterations have been investigated in the context of adenomyosis with the goal of discerning contributing genetic factors. Comparative genomic hybridization (CGH) analysis of frozen tissues from 25 cases of pathologically-proven adenomyosis did not detect recurrent chromosomal gains or losses, suggesting that large chromosomal changes may be rare [90].

Further, research by Chao et al. suggests a strong link between chromosomal instability and the pathogenesis of adenomyosis [91]. A whole exome and RNA sequencing study identified polyploidy (whole genome doubling, 4n) in 19 out 57 human adenomyosis tissue biopsies, which correlated with early disease onset and higher sensitivity to estrogen treatment [91].

Alternatively, a polyploid karyotype may suggest a metaplastic potential of our adenomyosis-derived cells. Metaplasia of displaced embryonic Müllerian remnants or stem cells is one of the theories of adenomyosis development [73, 92]. Karyotype abnormalities, including aneuploidy and non-random chromosomal breaks, are also found in adenomyosis patient-derived endometrial mesenchymal stem cells [93]. Interestingly, established herein tAEC21 cell line expresses cytokeratin (KRT)-5 (**Figure 1B**), a reported progenitor or stem cell marker, particularly in glandular reproductive and mammary tissues orchestrating cell maturation into luminal or myoepithelium [94, 95]. Immunohistochemical analysis by Stefansson et al. demonstrates low, focal expression of KRT5 in normal endometrium as opposed to its enhanced presence in endometrial hyperplasia and squamous metaplasia, which in turn is linked to endometrial epithelial malignancies [96, 97]. KRT5 positivity is associated with loss of E-cadherin expression, as well as suppression of ESR and PGR, which is consistent with our observations in tAEC21 [98, 99]. Furthermore, in reproductive cancers, KRT5+ cell percentage increases following cisplatin treatment, suggesting chemotherapy-induced enrichment of cancer stem cell population [94].

Altered response to high cytokine pressure contributes to adenomyosis symptomatology and progression and can be a promising therapeutic intervention target. Consistent with the biological properties of the disease, tAEC21 cells exhibit a significant increase in the production of multiple immune response mediators upon treatment with TNF-α (**Figure 5**), a key pro-inflammatory cytokine implicated in adenomyosis [61]. This response is characterized by elevated expression of IL6 and CXCL8 (IL8), cytokines known to amplify inflammatory signals and recruit immune cells, both of which are found at higher levels in both eutopic and ectopic endometrial tissues of adenomyosis patients [61, 100–102]. TNF-α stimulation of TAEC21 cells also led to a marked upregulation of CCL2, a chemokine critical for monocyte recruitment and subsequent macrophage infiltration, which is also elevated in adenomyosis tissues [61, 100, 101]. Macrophages are highly enriched in the eutopic and ectopic endometrium of adenomyosis patients, and their dysregulated activity, largely attributed to M2 polarization, facilitates EMT of endometrial epithelium, enhances invasiveness of adenomyotic lesions, and contributes to disease-associated infertility through the disruption of immune tolerance and subsequent embryo implantation failure (reviewed in [103]). Finally, TNF-α-stimulated tAEC21 cells increased expression of MUC1, a glycoprotein that is normally expressed on apical surfaces of uterine epithelial cells forming a protective barrier, but depending on the context, may promote or suppress inflammation, result in adaptive anti-MUC1 immunity, and alter the state of immune tolerance necessary for successful implantation [104–107]. Observed findings represent the initial characterization of tAEC21 cell line, and future studies will investigate a broader cytokine/chemokine panel to establish this cell line’s inflammatory profile.

3D cell culture systems have become increasingly valuable in mimicking the natural architecture of tissues and organs, faithfully representing cell-cell and cell-matrix interactions, and providing more accurate drug responses than cell monolayers. In this work, we screened tAEC21 cells for their propensity to self-organize and interact with endometrial stromal fibroblasts, confirming the spontaneous formation of physiologically relevant 3D spheroids with an epithelial tAEC21 shell surrounding a stromal core (**Figure 6**). Our observations mirrored results by Song et al. who presented similar spatiotemporal organization and epithelial-stromal interaction between epithelial-like 12Z cells and uterine or endometriotic stromal cells in an elegant spheroid model for endometriotic lesions [108]. In the absence of established adenomyosis cell lines, in vitro adenomyosis tissue 3D modeling relies on patient-derived organoid technology, often coupled with synthetic biomaterial matrices and microfluidic platforms (comprehensively reviewed by Gnecco and colleagues) [109]. Our described tAEC21/THESC epithelial/stromal spheroid model presents a valuable alternative to adenomyosis-derived organoids, offering greater simplicity, faster generation, and cost-efficiency without the need for post-hysterectomy tissue explants and artificial scaffolds. Given our past experience with the model, we predict easy manipulation of spheroid size and cellular complexity by incorporating various cell subpopulations, as well as straightforward drug testing in the future.

Immunolocalization studies report strong nuclear ESR expression in human adenomyotic glandular epithelium and stroma with varying levels across menstrual cycle phases [110, 111]. Progesterone receptor (PGR) expression reports are controversial. Some works confirm its nuclear-cytoplasmic presence in glandular epithelial and stromal cells of human adenomyosis tissue similar to normal myometrium [110]. Other studies show loss of PGR expression in adenomyotic samples, especially stroma basalis and myometrium [111]. Our initial assessment of 2D-cultured tAEC21 hormone receptor status showed undetectable levels of both ESR1 (ERα) and PGR via immunostaining. Importantly, our work showed gain of ERα expression by tAEC21 under 3D (co)culture conditions only **(Figure 4A**; **Figure 7**). This observation suggests that 3D context of cells fosters spatial cell-cell interactions and activates signaling pathways conducive to ESR expression [112, 113]. The lack of PGR expression in tAEC21 cells was maintained under both 2D and 3D culture conditions **(Figure 4A**; **Figure 7**). Interestingly, while not explicitly shown in adenomyosis, high Ncad expression negatively correlates with PGR expression in endometriotic ectopic lesions and endometriotic 12z cell line [46]. Furthermore, in a tamoxifen-induced adenomyosis mouse model, progression of ectopic lesions and an increase of EMT markers coincided with progressive loss of PGR in immunostained adenomyotic glandular epithelium [85]. Future detailed profiling of all hormone receptor subtypes is required.

In summary, our novel endometrial-like epithelial cell line derived from a patient with pathologically confirmed adenomyosis and a history of endometriosis exhibits characteristics of invasiveness, karyotypic abnormalities, accurately mimics immune responses observed in vivo, faithfully recapitulates physiologically relevant spatial rearrangement with supporting stroma in a heterotypic epithelial/stromal 3D spheroid model, and bears a conditional ESR+/- and a consistent PGR-status. Adequate growth rate in multiple commonly used purchasable media, supplemented with standard or charcoal/dextran-stripped FBS, makes tAEC21 cell line a flexible, inexhaustible, time- and cost-effective in vitro model that reduces media-related experimental constraints and removes ethical concerns associated with human and animal tissue use. It can serve as a valuable tool for exploring the underlying mechanisms of adenomyosis/endometriosis and associated complications, as well as testing new diagnostic and treatment strategies.

## ACKNOWLEDGMENTS

We thank Dr. Hari Nakshatri (Indiana University) for technical support and providing a pLXSN-hTERT vector. This work was supported in part by the National Institutes of Health (NIH), Eunice Kennedy Shriver National Institute of Child Health and Human Development (NICHD) (R01HD109707 and R21HD102653-01A1) to S.M.H and Y.K. and National Cancer Institute (R25CA269039-01A1) to S.M.H. and H.D.Q. We thank the Biospecimen Collection and Banking Core (BC^2^) at the Indiana University Simon Comprehensive Cancer Center (IUSCCC) who provided human tissue procurement service in support of this study. We thank contributors, including Indiana University, who collected samples and/or data used in this study, as well as study participants whose help and participation made this work possible. This work was supported by the Immunohistochemistry Research Core; the Histology Core of the Indiana Center for Musculoskeletal Health at Indiana University School of Medicine, and the Indiana Clinical Translational Sciences Institute (CTSI). We thank the IUSCCC Flow Cytometry Core for their outstanding technical support. The IUSCCC Flow Cytometry Core is funded in part by NIH, NCI grant P30 CA082709. The graphical abstract was created in https://BioRender.com

## AUTHOR CONTRIBUTIONS STATEMENT

Conceptualization, YK, SMH; Methodology, YK, SMH; Investigation, YK, JLK, HDQ; Formal Analysis, YK; Writing - Original Draft, YK; Writing - Review and Editing, YK, SMH; Visualization, YK, JLK, HDQ; Project Administration, YK; Supervision, SMH; Funding Acquisition, YK, HDQ, SMH. All authors have read and approved the final submitted version of the manuscript.

---

**Supplemental Figure S1.**
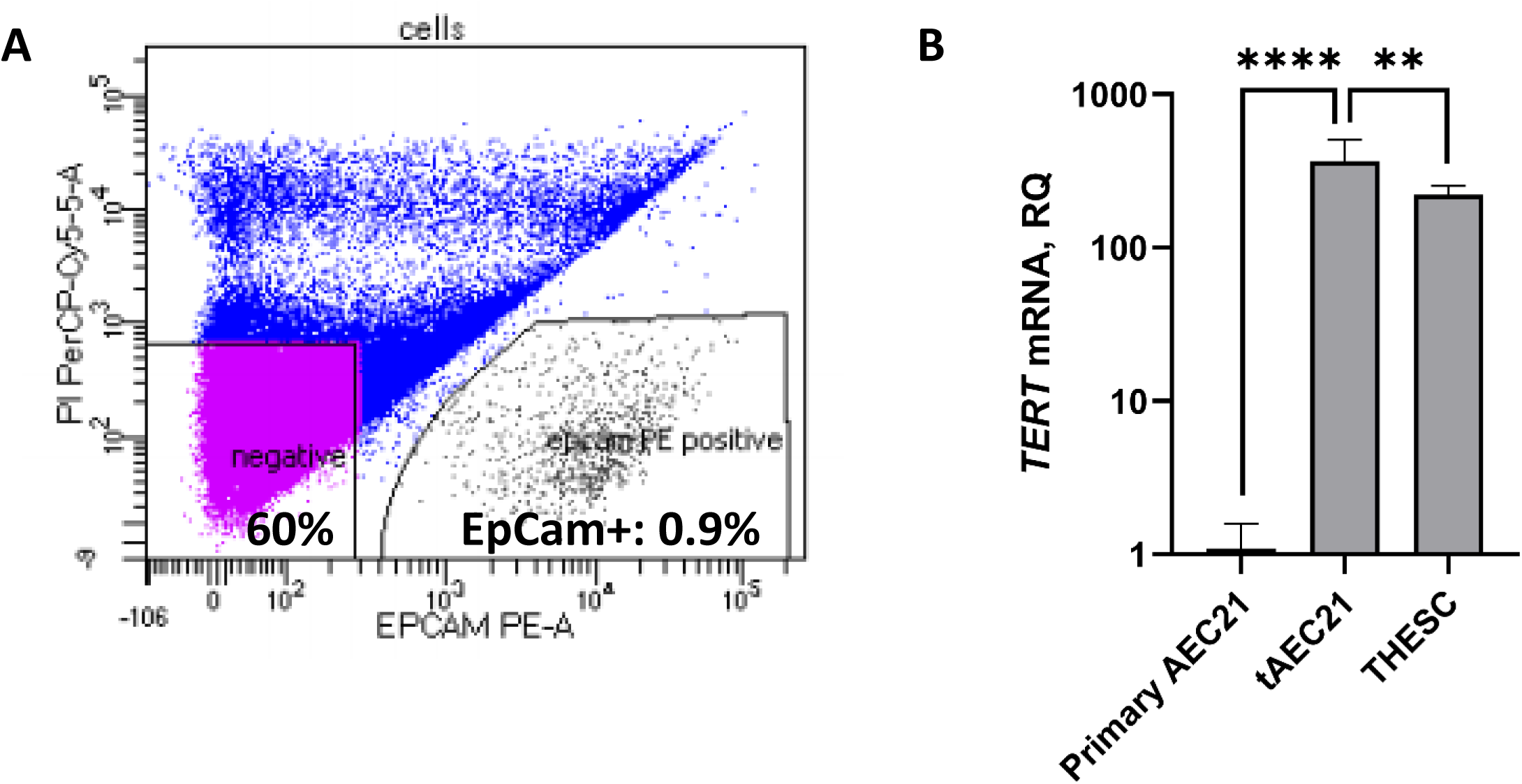
A) Human primary endometrial tissue cells from a uterus with adenomyosis were tagged with EpCam-PE antibody (#130-110-999, Miltenyi Biotec), and counterstained with propidium iodide (PI; Life Technologies, #PI304MP) for exclusion of dead cells; cells were sorted into EpCam+ (epithelial) and EpCam-(stromal) subpopulations by BD FACSAria Fusion cell sorter and BD FACSDiva Software. B) Primary AEC21 cells, hTERT-immortalized tAEC21, and established THESC cells (hTERT+ positive control) were cultured as monolayers for 24h; RNA was extracted and processed for qPCR. Data are presented as relative quantification (RQ) of human telomerase reverse transcriptase (*TERT*) expression in immortalized cells compared to primary culture, normalized to β-actin endogenous control (mean ± SEM, n≥3); **, p<0.01, ****, p<0.0001, unpaired two-tailed Student’s t-test. Statistical analysis was performed in GraphPad Prism (version 9.2.0).

**Supplemental Figure S2.**
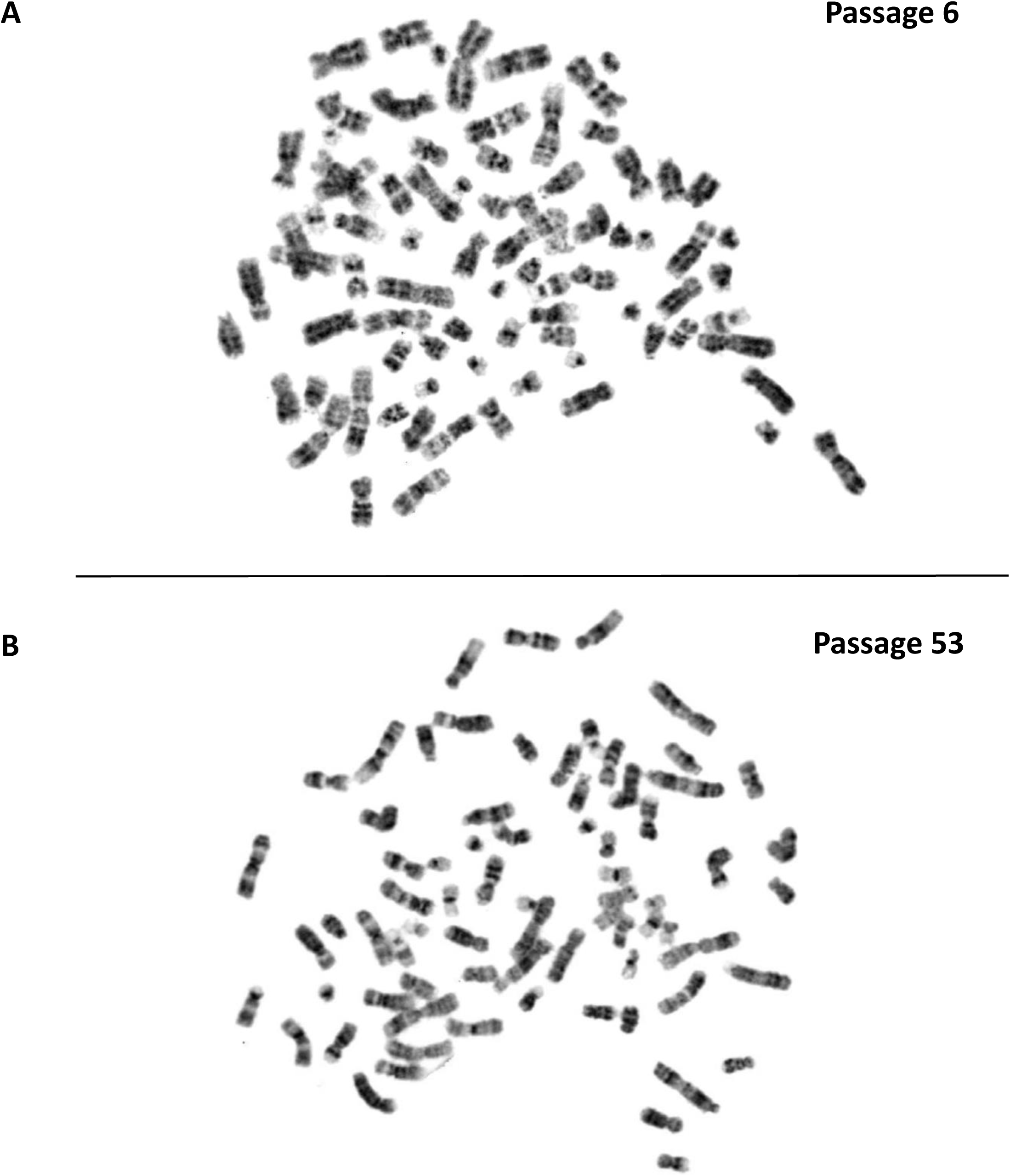
Simple G-banded metaphase chromosome cytogenetic analysis of tAEC21 cells at early and late passages. (A) Spread #9 of tAEC21 at passage 6 – 81 chromosomes with multiple rearrangements; (B) Spread #7 of tAEC21 at passage 53 – 74 chromosomes with multiple rearrangements.

**Supplemental Figure S3.**
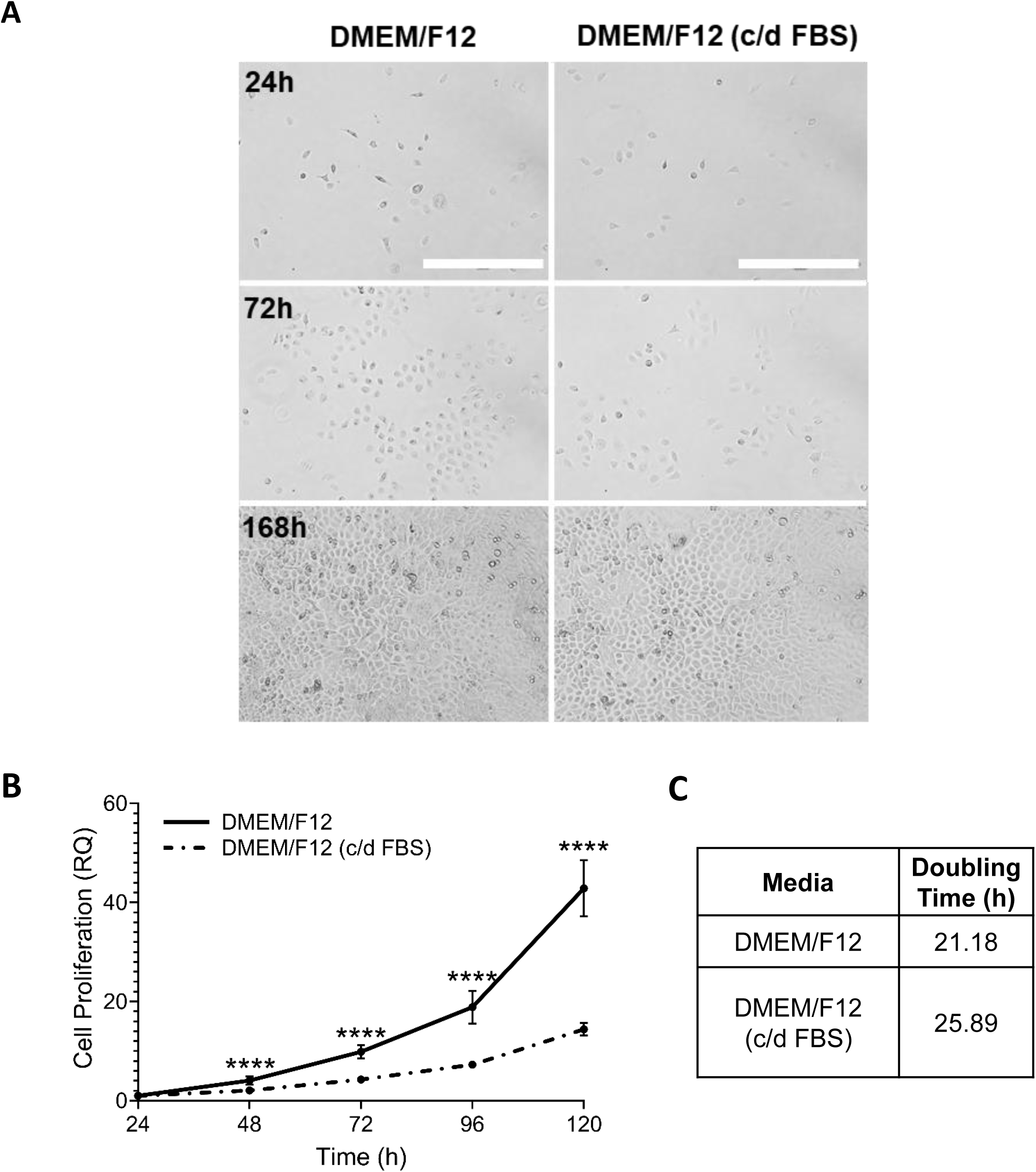
A) tAEC21 cells were seeded (25,000 cells/well) in a 6-well plate in different cell culture media (see Suppl. Table 1). Cell growth/morphology was monitored for 168 h (7 days) with daily fresh medium exchange (n=3). Images were taken using AMG EVOS FL imaging system; scale bar: 400 μm. B) tAEC21 were plated (500 cells/well) in a 96-well plate and subjected to MTS proliferation assay. Data are presented as relative quantification (RQ) with absorbance values normalized to 24 h post-seeding. Statistical analysis (one-way ANOVA with Tukey’s multiple comparison; ****, p < 0.0001; n=3) and C) cell doubling time acquisition (nonlinear fit) were performed in GraphPad Prism version 9.2.0.

